# Subthalamic Nucleus Neurons Differentially Encode Early and Late Aspects of Speech Production

**DOI:** 10.1101/227793

**Authors:** WJ Lipski, A Alhourani, T Pirnia, PW Jones, C Dastolfo-Hromack, LB Helou, DJ Crammond, S Shaiman, MW Dickey, LL Holt, RS Turner, JA Fiez, RM Richardson

## Abstract

Basal ganglia-thalamocortical loops mediate all motor behavior, yet little detail is known about the role of basal ganglia nuclei in speech production. Using intracranial recording during deep brain stimulation surgery, we tested the hypothesis that the firing rate of subthalamic nucleus neurons is modulated in response to both planning and motor execution aspects of speech. Nearly half of 79 recorded units exhibited firing rate modulation, during a syllable reading task administered in 12 subjects. Trial-to-trial timing of changes in subthalamic neuronal activity, relative to cue onset versus production onset, revealed that locking to cue presentation was associated more with units that decreased firing rate, while locking to speech onset was associated more with units that increased firing rate. These uniquely acquired data indicate that subthalamic activity is dynamic during the production of speech, reflecting temporally-dependent inhibition and excitation of separate populations of subthalamic neurons.

## INTRODUCTION

Producing speech is the most complex of human motor behaviors, requiring dynamic interaction between multiple brain regions. The segregated loop organization of basal ganglia-thalamocortical circuits suggests that the basal ganglia, including the subthalamic nucleus (STN), play a critical role in speech production. This concept is supported additionally by observations that impairments in speech production are common features of basal ganglia-associated degenerative disorders including Parkinson’s disease, and that other disorders in speech production (e.g., stuttering) are associated with abnormalities in basal ganglia activity (Alm, 2004; Giraud et al., 2008; Toyomura et al., 2015). Additionally, an extensive body of work in song birds implicate bird-homologues of the basal ganglia (Doupe and Kuhl, 1999), including a homologue of the STN, in the learning and production of vocalizations (Jiao et al., 2000). Many prominent models of speech production nonetheless virtually ignore the basal ganglia (Hickok, 2012), as few studies have examined speech-related neural activity in these subcortical nuclei directly (Ziegler and Ackermann, 2017).

Electrophysiological recordings obtained during the implantation of leads for deep brain stimulation (DBS) represent the only clinically-indicated opportunity to measure neural activity directly from the basal ganglia in awake, behaving human subjects. Previous reports of STN unit activity, however, have been limited to only a single preliminary, qualitative analysis of speech production-related changes in STN firing rates (Watson and Montgomery, 2006). Thus, recording from STN neurons during speech production is a unique opportunity to test hypotheses about the role of this region in the control of complex motor function, where the basal ganglia have alternately been hypothesized to participate in action selection, movement gain and motor learning (Desmurget and Turner, 2010).

To begin to define the role of the STN in speech production more clearly, we established an intraoperative protocol for microelectrode recording during a task that required subjects to read aloud single syllables displayed on a computer screen. We then examined trial-to-trial timing of changes in STN unit activity relative to either the visual presentation of single syllables or to the onset of speech production. Such time-locking is considered as evidence for an underlying functional linkage between the behavioral event and the linked neural discharge (Seal and Commenges, 1985; Anderson and Turner, 1991; DiCarlo and Maunsell, 2005). Our results suggest that aspects of speech production are encoded in the STN through the inhibition and excitation of functionally segregated neurons.

## RESULTS

Subject demographics are summarized in Table 1. Twelve subjects each performed between 1 and 4 blocks of 60 trials during single unit recording sessions (median 2.5 blocks, mean 160 trials). An average 6.5 ± 1.9% of trials were excluded from analysis due to incorrect responses. Across subjects, the mean latency to the onset of a production was 1.10 ± 0.31s, and the mean duration of speech was 0.605 ± 0.175s. A subject’s mean production latency correlated significantly with the subject’s speech UPDRS sub-score (Spearman rho = 0.72, p = 0.02). This correlation failed to reach significance for speech duration (Spearman rho = −0.09, p = 0.8) or the fraction of trials with incorrect responses (Spearman rho = −0.62, p = 0.06). The subjects’ total UPDRS score was not correlated with any of these task measures (production latency: Spearman rho = 0.22, p = 0.5; speech duration: Spearman rho = −0.23, p = 0.5; percent correct: Spearman rho = 0.22, p = 0.5).

**Table 1.** Subject Characteristics.

| Subject | Age | sex | UPDRS off score |  | # sessions | mean production latency (s) | production latency S.E. (s) | mean speech duration (s) | speech duration S.E. (s) | % correct trials |
| --- | --- | --- | --- | --- | --- | --- | --- | --- | --- | --- |
|  |  |  | speech | total |  |  |  |  |  |  |
| 1 | 60 | male | NR | 53.0 | 2 | 1.50 | 0.03 | 0.777 | 0.005 | 89% |
| 2 | 68 | male | 1 | 47.0 | 4 | 1.01 | 0.14 | 0.578 | 0.016 | 95% |
| 3 | 47 | female | NR | NR | 3 | 0.91 | 0.03 | 0.623 | 0.010 | 100% |
| 4 | 60 | male | 0 | 31.0 | 2 | 0.77 | 0.04 | 0.642 | 0.020 | 98% |
| 5 | 68 | male | 1 | 50.0 | 2 | 0.75 | 0.00 | 0.592 | 0.039 | 97% |
| 6 | 56 | male | 1 | 45.5 | 2 | 0.86 | 0.00 | 0.467 | 0.009 | 98% |
| 7 | 82 | male | 2 | 36.0 | 3 | 1.99 | 0.13 | 0.442 | 0.007 | 76% |
| 8 | 66 | male | 0 | 45.5 | 4 | 0.67 | 0.02 | 0.610 | 0.007 | 97% |
| 9 | 66 | male | 2 | 45.0 | 2 | 1.13 | 0.02 | 0.745 | 0.023 | 93% |
| 10 | 71 | female | 1 | 24.0 | 3 | 1.26 | 0.06 | 0.796 | 0.016 | 86% |
| 11 | 77 | male | 1 | 27.0 | 2 | 1.33 | 0.25 | 0.539 | 0.012 | 96% |
| 12 | 59 | male | 1 | 40.0 | 3 | 1.04 | 0.10 | 0.452 | 0.004 | 96% |

NR = not recorded, S.E. = standard error

A total of 45 neuronal recordings met the criteria for A-sorts (22 single-unit, 23 multi-unit recordings). Thirty-four additional recordings met criteria for B-sorts (3 single-unit, 31 multi-unit recordings). Similar to firing rate data reported previously from the human STN (Rodriguez-Oroz et al., 2001; Abosch et al., 2002; Starr et al., 2003; Theodosopoulos et al., 2003; Romanelli et al., 2004; Schrock et al., 2009), The mean baseline firing rates were not significantly different between A- and B-sorts (21.8 ± 3.2 spikes/s vs. 27.3 ± 7.1 spikes/s; mean ± standard error, p = 0.55, unpaired t-test). These groups were combined subsequently for analysis.

Overall, a high percentage of units demonstrated a speech-related change in firing (Table 2). 22 units exhibited significant increases in firing rate, 13 units showed significant decreases, and 7 units showed a mixed increase/decrease response during the production epoch. Figure 1A-C shows examples of these three unit response categories. While there was an overall significant difference in the proportions of neurons in the four response categories (increase, decrease, mixed, and non-response, χ^2^ = 25.8608, p = 1.02 × 10^−5^), there was no significant difference between the proportion of increase-type and decrease-type units (χ^2^ = 2.3 p = 0.13). The prevalence of speech-responsive units did not relate to the subjects’ symptom severity, and the proportion of units recorded from each subject that showed increase, decrease, cue-locking or speech-locking response types was not correlated with the speech sub-score or total UPDRS score (Table 3). Increase- and decrease-type single-units were not differentiated statistically by baseline firing rates (increase-type firing rate = 20.6 ± 6.4 spikes/s; decrease-type firing rate = 34.4 ± 10.5 spikes/s; p = 0.27, unpaired t-test). The mean latency of neuronal responses (defined as onset of the first significant change relative to production onset, see Figure 1D-F) also was similar between increase and decrease response types (-0.23 ± 0.07s and −0.20 ± 0.14s, respectively, p = 0.87, unpaired t-test).

**Table 2.** Number of units recorded in each response category.

|  | Total | No response | Increase | Decrease | Mixed |
| --- | --- | --- | --- | --- | --- |
| A-sorts | 45 | 19 | 13 | 10 | 3 |
| B sorts | 34 | 18 | 9 | 3 | 4 |
| **total** | **79** | **37** | **22** | **13** | **7** |

**Table 4.** Subjet symptom severity is not correlated with unit speech response types.

| Correlation with Speech UPDRS | proportion of units by response type |  |  |  |
| --- | --- | --- | --- | --- |
|  | increase-type | decrease-type | speech-locked | cue-locked |
| Spearman $\rho$ | 0.28 | 0.08 | 0.39 | 0.45 |
| p-value | 0.44 | 0.82 | 0.27 | 0.20 |
| Correlation with Total UPDRS | proportion of units by response type |  |  |  |
|  | increase-type | decrease-type | speech-locked | cue-locked |
| Spearman $\rho$ | 0.04 | -0.03 | -0.51 | -0.30 |
| p-value | 0.92 | 0.92 | 0.11 | 0.36 |

**Figure 1.**
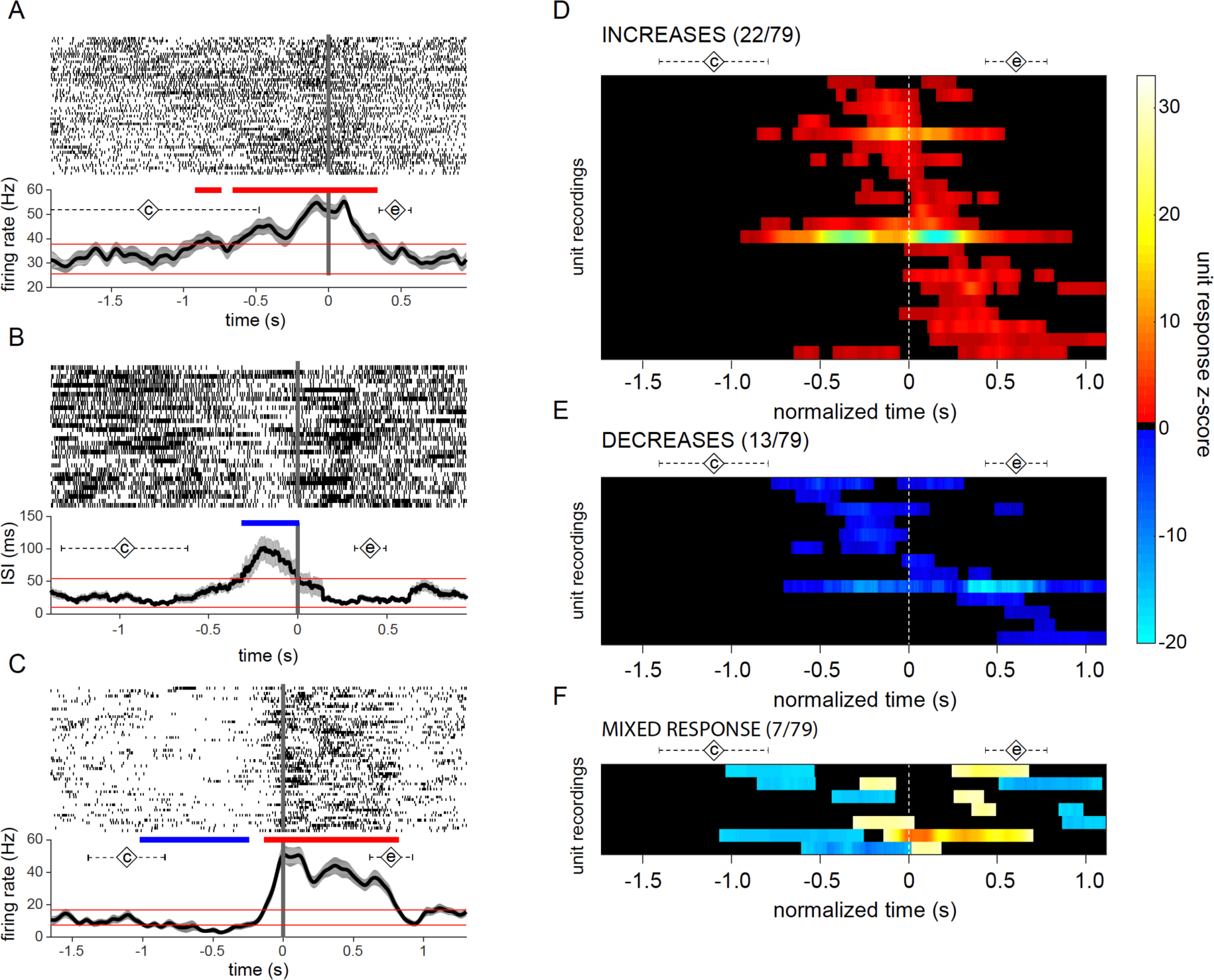
STN neuronal firing is modulated during speech. Examples of neuronal responses during speech showing (A) increases, (B) decreases and (C) mixed responses in firing rate, aligned to production onset (t=0). Spike rasters across trials are shown on top in panels A-C, and mean firing rate (A, C) or mean inter-spike interval (ISI; B) is shown on the bottom. Diamonds labeled with a “c” indicate mean time of cue presentation; diamonds labeled with an “e” indicate mean speech end; dashed error bars indicate the corresponding standard deviations. (D-F) Raster plots illustrating the timing of firing rate responses across the population of unit recordings. Each row represents a unit’s significant changes relative to baseline, during a time segment surrounding production onset. The time scale is normalized across units from 0.5 s before the mean cue onset until 0.5 s after the mean end of speech. Diamonds labeled with a “c” indicate subject mean time of cue presentation; diamonds labeled with an “e” indicate subject mean speech end; dashed error bars indicate the corresponding standard deviations.

Response types were observed to be differentially associated with speech onset- and cue- locking. Among 29 units with significant increases in firing rate during the production epoch, the responses were preferentially time-locked to production (41%), with a minority time-locked to cue onset (7%) or to both cue and production onset (7%) (Figure 2). In contrast, among 20 units with significant decreases in firing rate, 40% were time-locked to cue onset, while only 15% had responses time-locked to production onset, and no responses were time-locked to both cue and production onset (Figure 3). A Chi-square test was used to verify that increase-type neural responses were more likely to be time-locked to the production onset than were decreases (χ^2^ = 3.89, p = 0.049), whereas decrease-type responses were more likely to be time-locked to cue onset (χ^2^ = 7.99, p = 0.0047). The mean latency of neuronal responses (defined as the as the mean neuronal response latency across trials) was shorter for cue-locked responses (0.76 ± 0.12 s) than for speech-locked responses (1.08 ± 0.08 s; p = 0.039, unpaired t-test). The mean neuronal response to production interval (defined as the mean neuronal response to production interval across trials) was also greater in magnitude for cue-locked responses (-0.48 ± 0.12 s) than for speech-locked responses (-0.15 ± 0.6 s; p = 0.011, unpaired t-test).

**Figure 2.**
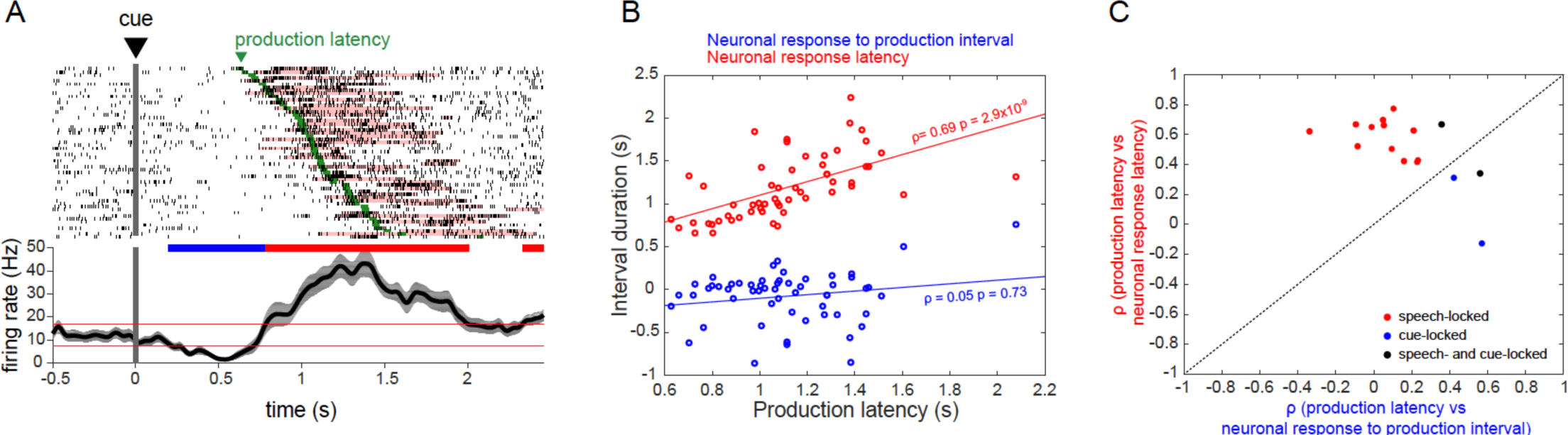
STN neuronal firing increases are primarily speech-locked. (A) An example of a neuron whose firing rate increase is locked to production onset. Spike raster (top) and mean firing rate (bottom) aligned to cue presentation. Significant spike bursts are shaded for each trial according to their Poisson Surprise index. Trials are sorted by speech production latency; speech production onset for each trial is indicated in green. (B) The time interval between cue presentation and burst onset (neuronal response latency) and between burst onset and production onset (neuronal response to production interval) for each trial is correlated against production latency. (C) Summary of correlation analyses for all unit recordings with increase-type responses, showing units locked to production onset (red circles), locked to cue presentation (blue circles), and locked to both cue and production onset.

**Figure 3.**
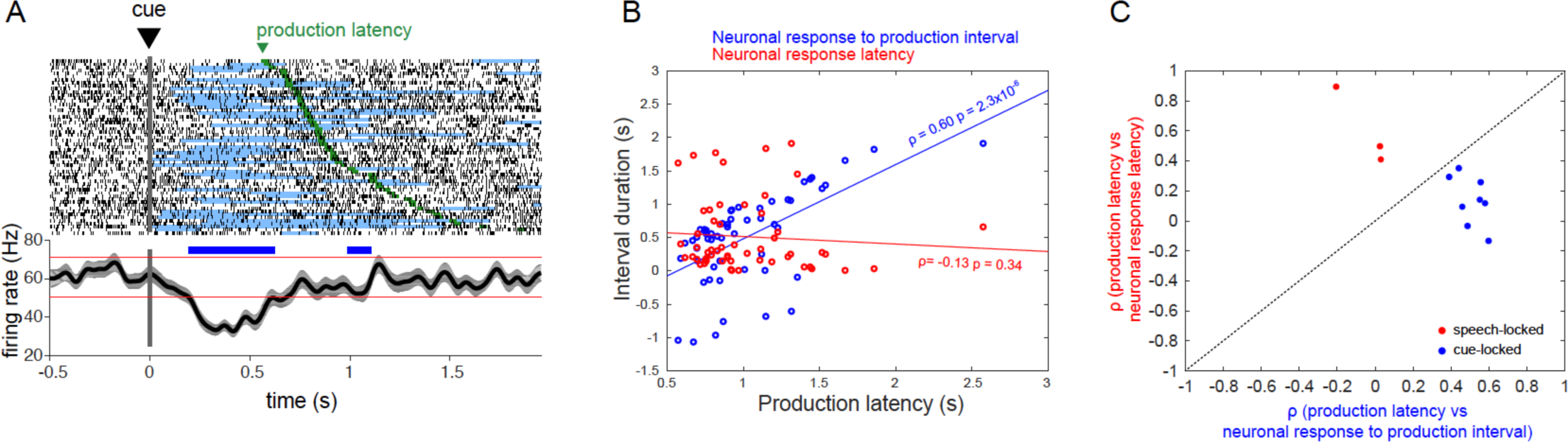
STN neuronal firing decreases are primarily cue-locked. (A) An example of a neuron whose firing rate decrease is locked to cue presentation. Spike raster (top) and mean firing rate (bottom) aligned to cue presentation. Significant decreases in firing rate (pauses) are shaded for each trial according to their Poisson Surprise index. Trials are sorted by speech production latency; speech production onset for each trial is indicated in green. (B) The time interval between cue presentation and pause onset (neuronal response latency) and between pause onset and production onset (neuronal response to production interval) for each trial is correlated against production latency. (C) Summary of correlation analyses for all unit recordings with inhibitory responses, showing 3/20 units were locked to production onset, 8/20 units showed locking to cue presentation, and none locked to both cue presentation and production onset.

Encoding of speech duration was not prevalent in recorded STN units. The duration of the neural response had a significant correlation with the duration of speech production (Spearman correlation, p < 0.05) in only 2 of 29 units with increase-type responses, and in only 1 of 20 neurons with decrease-type responses. These proportions were not significantly different from zero (Fisher’s exact test, p = 0.49 for increase-type responses, p = 1.0 for decrease-type responses), indicating that they are too small to be estimated statistically from this experiment.

We did not find evidence for topographical organization of response types. Unit recording locations were analyzed based on the recording trajectory (center, 23 units; posterior, 29 units; or medial, 27 units), and the recording depth, relative to the microelectrode recording-defined boundaries of the STN within each trajectory. There was no significant difference in STN recording depth between speech response types (excitatory, inhibitory, mixed, no response; see Figure 4) in any of the recording trajectories (Kruskal-Wallis test, central trajectory χ^2^ = 7.2, p = 0.066; posterior trajectory χ^2^ = 6.2, p = 0.10; medial trajectory χ^2^ = 7.2, p = 0.066). There was also no significant difference in STN recording depth between locking response types (production onset-locked, cue-locked, locked to both events, no locking) in any of the recording trajectories (Kruskal-Wallis test, central trajectory χ^2^ = 5.6, p = 0.13; posterior trajectory χ^2^ = 3.9, p = 0.27; medial trajectory χ^2^ = 2.1, p = 0.35). Collapsing the recording depths across trajectories did not result in a significant difference in depth between response types.

**Figure 4.**
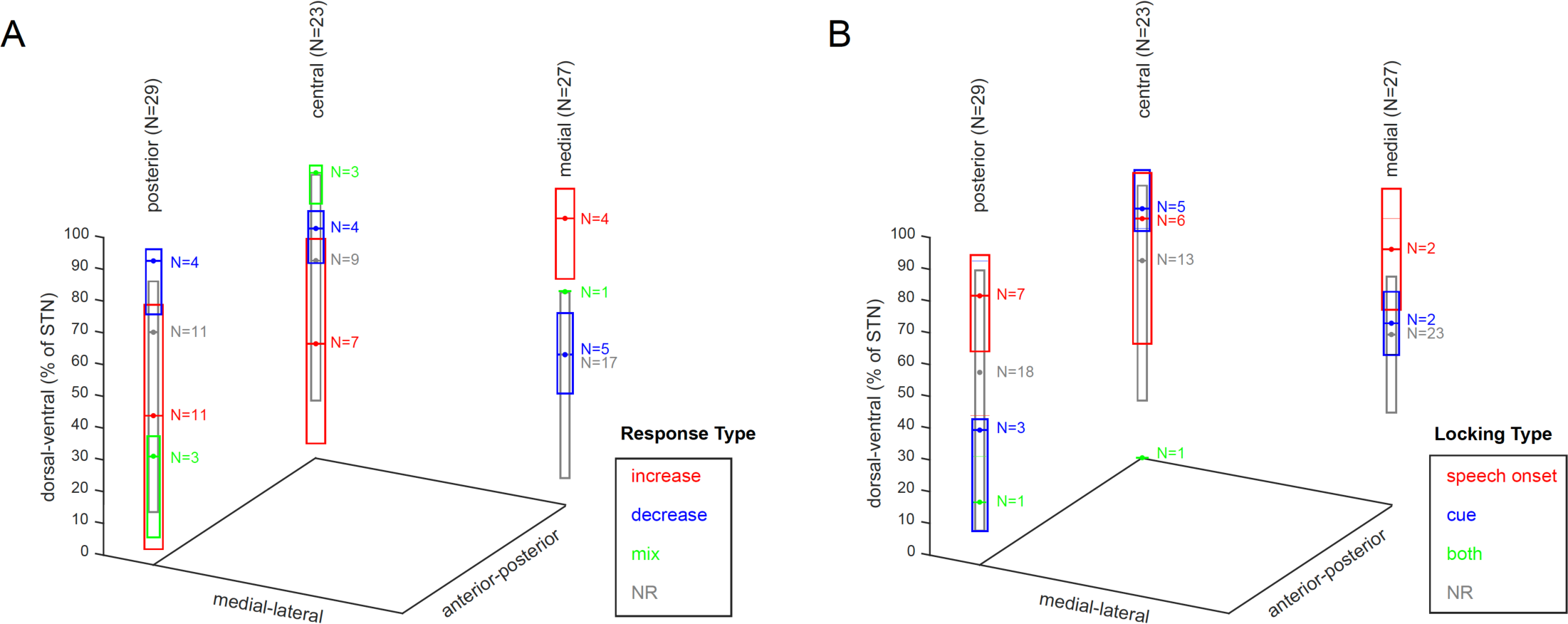
Anatomical distribution of speech responses in the STN. Units locations are represented according to the recording trajectory and recording depth relative to electrophysiology-defined STN boundaries (0% corresponds to the ventral STN border and 100% corresponds to the dorsal STN border. Box plots represent the median and inter-quartile range of recording depths within each response category.

## DISCUSSION

We found that both phasic increases and decreases in the discharge rate of STN neurons accompany the production of speech. In this study, subjects read aloud syllables presented on a computer screen, a behavioral paradigm that requires a series of neural events beginning from processing the visual cue to activating motor commands for the vocal organ. Neural events that occur early in this series, such as processing of the visual cue and forming a phonological plan, might be expected to be time-locked to cue presentation. Events that occur later in the series, such as forming and executing the motor speech plan, might be expected to be time-locked to speech output. We showed that decrease-type responses are predominantly locked to cue presentation and increase-type STN responses are predominantly locked to the onset of speech. These findings suggest that STN inhibition may be associated with early, cognitive aspects of speech production, while STN excitation may be associated with later, motor aspects of speech production.

The extent to which speech-related activity in the STN may reflect lower-order movement-related activity, akin to results from studies involving simple limb movements, versus higher order functions has important implications. Although kinematic aspects of speech production often improve following DBS (Pinto et al., 2004; De Gaspari et al., 2006; Parsons et al., 2006; Mikos et al., 2011), a decrease in verbal fluency is the most common cognitive side effect of STN-DBS, with specific deficits in lexical and grammatical processing having been observed, albeit inconsistently across studies (Phillips et al., 2012). The observation of increases in firing rates associated with speech onset are expected, in the context of previous studies of limb movement-related activity. In STN recordings from both human subjects and non-human primates, firing rate increases comprise 75-93% of movement-related responses during active and passive limb movements (Wichmann et al., 1994; Rodriguez-Oroz et al., 2001; Abosch et al., 2002; Starr et al., 2003; Theodosopoulos et al., 2003; Romanelli et al., 2004; Schrock et al., 2009). We found that nearly half of increase-type responses in our study were locked to the onset of speech, indicating that motor aspects of speech production are encoded in STN activity. A significantly smaller proportion of increase-type responses was locked to cue presentation (7%) and to both cue presentation and speech production onset (7%), with remaining responses not clearly associated with either event.

In contrast, we observed that early stages of speech production may involve the inhibition of STN neurons. We found that a large proportion (40%) of decrease-type responses were locked to cue presentation, with cue-locked responses occurred at significantly lower latencies relative to cue presentation, compared with speech-locked responses. A smaller proportion (15%) of decrease-type responses were locked to speech production onset, with remaining responses not clearly associated with either event. Altought minority populations of neurons with movement-related firing-rate decreases have been reported previously (Wichmann et al., 1994; Schrock et al., 2009; Lipski et al., 2017), and active movements have been associated with a higher proportion of decrease-type responses in the STN (Lipski et al., 2017), it is remarkable that such a high percentage of decrease-type responses were observed in the present study. Interestingly, and in contrast to our results, a marked reduction of STN activity was reported to be associated with the onset of speech production in the only previous report of STN unit activity recorded during speech production (Watson and Montgomery, 2006), although that study was largely descriptive in nature, limiting comparisons to our data. Although other investigators have shown correlations of STN single unit firing rates and rhythms to premotor functions, such as the encoding of difficulty level of a choice task (Zaghloul et al., 2012; Zavala et al., 2016), cue-locked decreases in firing were not reported. Our data do suggest that, in comparison to limb movement, speech may involve a different balance of activation and suppression in the STN, and that modulation of this balance may occur at the single neuron level prior to speech onset.

The task-related activity of STN neurons, such as that observed here, are presumed to function within the basal ganglia thalamocortical circuit primarily by way of glutamatergic inputs to the GABAergic output neurons of the globus pallidus internus and substantia nigra pars reticulata. The firing rate model of basal ganglia function posits that increases in STN activity may have a suppressive effect on basal ganglia-recipient circuits while decreases may be facilitatory. This balance of basal ganglia-mediated activation and suppression has been understood most frequently in terms of either selecting and focusing motor actions (Mink, 1996; Redgrave et al., 2010), or modulating their gain over time (Alexander and Crutcher, 1990a; Nambu et al., 2000, 2002; Nambu, 2005; Turner and Desmurget, 2010; Thura and Cisek, 2017). Proponents of an action selection hypothesis have proposed that the STN participates in a response inhibition function to reduce premature action when multiple competing responses are possible (Frank, 2006). Our findings of suppressed STN firing locked to speech cues and increased STN firing locked to speech production, however, are not consistent with action selection-related functions of the STN. Similarly, Zeigler and Ackerman (Ziegler and Ackermann, 2017) recently compiled extensive evidence in support of the idea that, for well-learned adult speech, basal ganglia circuits play key roles in the emotional/motivational modulation of speech (i.e., in prosody) but not in the selection and sequencing of articulatory gestures.

Speech-related phasic increases in the STN likely are a result of excitatory inputs, and decreases likely a result of inhibitory inputs. The major excitatory input into the STN comes from the neocortex via the basal ganglia hyper-direct pathway (Nambu et al., 2002) which forms glutamatergic synapses onto distal dendrites of STN projection neurons (Künzle, 1978; Romansky et al., 1979; Kitai and Deniau, 1981; Romansky and Usunoff, 1987). The primate STN receives direct projections from broadly distributed cortical areas including primary motor cortex, pre-motor cortex, supplementary motor area, dorsolateral prefrontal, anterior cingulate, and inferior frontal cortex (Afsharpour, 1985; Parent and Hazrati, 1995; Nambu et al., 1997; Haynes and Haber, 2013). A primary form of inhibitory drive arises from GABAergic projections to the STN from the external segment of the globus pallidus, via the indirect basal ganglia pathway (Nauta and Mehler, 1966; Romansky and Usunoff, 1987; Bell et al., 1995; Sato et al., 2000). Thus, it is possible that speech onset-locked increased firing rate responses (STN excitation) could be mediated via the hyper-direct pathway, while cue-locked inhibitory responses during speech could be mediated via the indirect pathway. These findings also can be interpreted in the context of the GODIVA model (Bohland et al., 2010) of speech production. This model posits a dual role for the basal ganglia, participating in two processes that may be correlated with cue presentation and speech production in our task: (1) a planning loop that is involved in generating a phonological sequence corresponding to the target word, and (2) a motor loop that releases the planned speech sounds for motor execution.

In summary, our results demonstrate that STN neurons comprise separate functional populations whose activity during speech production can be differentiated by the timing and direction of firing rate changes. The extent to which these functional groupings may be specific to speech versus common to complex motor function is an important question for future work, in light of conflicting theories of the role of the STN, and that of the basal ganglia as a whole, in motor behaviors. Our ongoing studies aim to examine the granularity of STN functional encoding in and to verify the specificity of these findings to speech production.

## METHODS

### Subjects

Subjects were 12 movement disorders patients (10 male) undergoing awake DBS surgery for Parkinson’s disease. Unified Parkinson’s disease rating scale (UPDRS) testing was administered by a neurologist within four months before DBS surgery. All subjects underwent overnight withdrawal from or a reduced dose of their dopaminergic medication prior to surgery. All participants provided written, informed consent in accordance with a protocol approved by the Institutional Review Board of the University of Pittsburgh (IRB Protocol # PRO13110420).

### Speech Task

The speech task was performed during pauses in the microelectrode recording portion of DBS lead implantation in which stable units were detected. Visual stimuli were created using Matlab software (MathWorks, Natick, MA) and Psychophysics Toolbox extensions (Brainard, 1997; Pelli, 1997; Kleiner et al, 2007). Subjects were asked to name a consonant-vowel-consonant (CVC) syllable presented in white text on an otherwise dark computer screen. Each trial was initiated manually by the experimenter, beginning with presentation of a green fixation cross at the center of the screen (0-250 ms), followed by a variable-time delay (500-1000 ms) during which the screen remained dark. At the end of the delay, text denoting a unique CVC syllable appeared on the screen and remained visible until the subject completed their naming response. A white fixation cross was displayed on the screen during the inter-trial interval (ITI; Figure 1A). Subjects were instructed to respond as soon as the word cue appeared.

### Electmphysiological recordings

Unit recordings were carried out using the Neuro-Omega recording system and Parylene insulated, microphonics-free tungsten microelectrodes (Alpha Omega, Nazareth, Izrael). Targeting and microelectrode recording (MER) were performed using standard clinical techniques for STN DBS (Starr et al., 2002). For each subject, two to three simultaneous microelectrode recording passes were made, using a center, posterior and medial trajectories spaced 2mm apart in a standard Ben-Gun xarray. Microelectrode signals were band-pass filtered at 0.075 Hz to 10 kHz and digitized at 44 kHz (NeuroOmega, Alpha Omega, Nazareth, Izrael).

### Audio recordings

Speech output was recorded using an omnidirectional microphone (8 subjects: Audio-Technica, Stow, OH; model ATR3350iS, frequency response 50-18,000 Hz; 4 subjects: Preosonus, Baton Rouge, LA, model PRM1 Precision Flat Frequency Mic, frequency response 20-20,000 Hz) oriented at an angle of approximately 45 degrees and a distance of approximately 8 cm to the subject’s mouth. In the four cases where the Preosonus PRM1 microphone was used, a Zoom H6 digital recorder was used to digitize the audio at 96 kHz. In all cases, the audio signal was split out to a Grapevine Neural Interface Processor, where it was digitized at 30 kHz. The audio signal was synchronized with the neural recordings and with visual cue events using digital pulses delivered via a USB data acquisition unit (Measurement Computing, Norton, MA, model USB-1208FS).

### Task performance

The audio signal was segmented into trials and responses were coded by a speech-language pathologist using a custom-designed graphical user interface implemented in MATLAB. The response epoch for each trial was defined to start at cue presentation and end at the start of the ITI. The audio signal within each response epoch was coded as follows: (1) production onset was identified, (2) production offset was identified, (3) the phonetic content was identified (Supp. Table 1). Only trials that met the following criteria were included for further analyses: (1) the subject’s entire response could clearly be identified within the response epoch; (2) the time from cue presentation to production onset (production latency) was less than the mean production latency (1.2 s) plus 3 standard deviations (0.93 s) for all subjects (threshold = 4.0 s); (3) the duration of the response (SpDur) was less than the mean production latency (0.60 s) plus 3 standard deviations (0.20 s) for all subjects (threshold = 1.19 s); (4) the subject’s response was a CVC or CV syllable and was composed of phonemes within the target set or the included mismatch set. Of 2,200 total trails, 150 (6.8%) were rejected from further analysis on the basis of these response criteria. In 11 of the rejected trails, no response was recorded. In 608 trials (139 of which were rejected), the response did not match the target.

### Spike sorting

Microelectrode recording data were imported into off-line spike-sorting software (Plexon, Dallas, TX) for discrimination of single- and multi-unit action potentials using principal components analysis. The results were graded according to the quality and stability of the spike sorting over the duration of the recording. An assignment of “A-sort” was given only to spike clusters that could be discriminated from background activity throughout the duration of a recording, and whose spikes were not strongly modulated by cardiac rate (see Figure 5C-F). A-sorts were further subdivided into single- and multi-unit subcategories. A cluster qualified as a single unit (SU) if: (1) the principal component cluster was clearly separated from other clusters associated with background activity and other units, (2) contained spike waveforms with a unimodal distribution in principal component space, and (3) displayed a refractory period of at least 3ms in its inter-spike interval distribution (Starr et al., 2003; Schrock et al., 2009). For some SU recordings, the location of the principal component cluster drifted gradually during the period of the recording, likely due to a shift of the brain relative to the electrode. Other A-sorts were classified as multi-unit (MU) recordings because the principle components cluster appeared to include waveforms from multiple units, forming multimodal principal component distributions that could not be clearly separated on short time scales, or that failed to obey the 3 ms refractory period in their inter-spike interval distribution. An assignment of “B-sort” was given to spike clusters that violated the above criteria due to presence of a non-uniform or rapid (5 second time scale) shift of the waveform cluster in principal component space, or due to incomplete separation of the spike cluster from the cluster associated with background noise.

**Figure 5.**
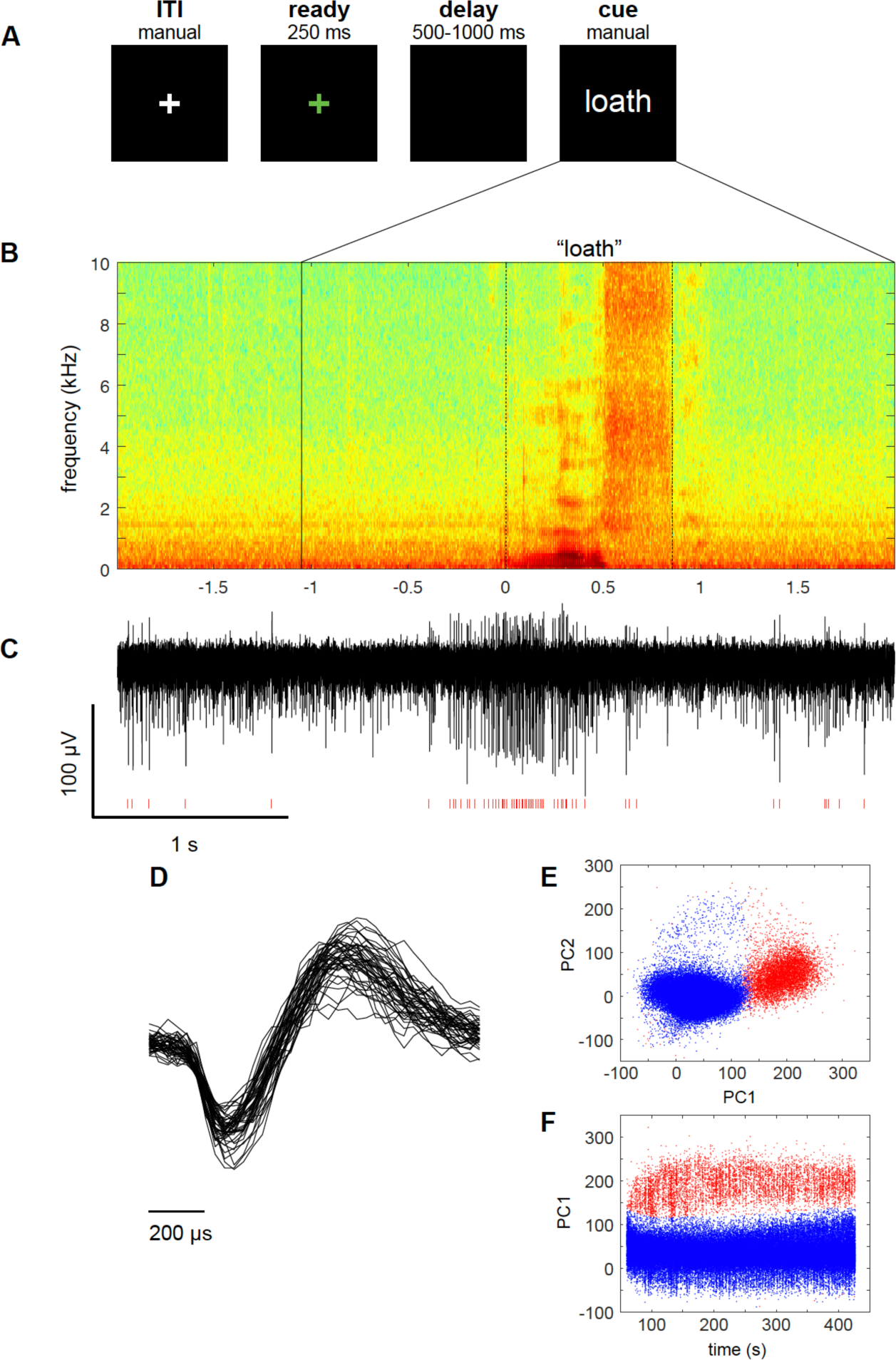
Speech task and representative spoken and neural responses. (A) Intraoperative syllable speech task. Subjects were asked to read aloud words presented on a computer screen. Each trial consisted of a sequence beginning with the fixation cross turning green for 250 ms, followed by a variable delay black screen (500-1000 ms), followed by a unique CVC syllable cue appearing on the screen until the response was recorded. A white fixation cross appeared during the inter-trial interval. (B) An example audio spectrogram time-aligned to the onset of a subject’s utterance of the syllable “loath.” The time (in s) of cue presentation is indicated by the solid vertical line, and the response onset and offsets are indicated by dotted lines. (C) A single unit recording from the subject’s STN, showing an increase in firing during speech. Red hash marks indicate timing of detected spike waveforms from the background activity. (D) Overlay of 50 spike waveforms from the single unit shown in (C). Scatterplots of the first two principal components (E; principal component1 and principal component2), as well as the first principal component and spike timestamp (F), showing clear separation of single unit spike waveforms (red) corresponding to the example shown in (C) from background (blue).

### STN unit activity during speech

We used two different estimates of unit activity to test for task-related changes in neuronal spike rate. We tested for task-related increases using a spike density function (SDF), which is a direct representation of a unit’s mean instantaneous firing rate. We tested for task-related decreases using a function that reflects a unit’s mean inter-spike interval (ISI), which scales with the reciprocal of instantaneous spike rate. This approach was chosen to avoid potential under-sensitivity for the detection of decreases in firing in SDFs due to floor effects (Alexander and Crutcher, 1990b). To construct an SDF function, spike time stamps were rounded to 1 ms. The resulting time series was then convolved with a Gaussian kernel (σ = 25 ms). The inter-spike interval (ISI) time series was computed from the 1 kHz binned time stamp time series by taking the value of the current ISI at each millisecond time point:

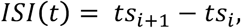

for *t* between *ts_i_* and *ts*_*i*+1_, where *ts* is the set of consecutive time stamps for that spike population.

Across-trial means of the SDF and ISI functions were constructed aligned on two epochs of interest: (1) from cue presentation to 0.5 s after the mean production onset for that session (aligned on cue presentation, termed the cue epoch), and (2) from the mean time of cue presentation to 0.5 s after production onset (aligned on production onset, termed the production epoch). A baseline period for each trial was defined as the 1 s portion of the ITI preceding cue presentation, and the trial-wise mean SDF and ISI functions during this epoch served as baselines against which the test epochs were compared. Baseline firing rates for each unit were defined as the mean of discharge rate during the baseline period across trials.

A unit was considered to have significantly elevated firing during a given epoch if the mean spike density within that test epoch exceeded a threshold level for at least 100ms. The threshold was defined as the upper 5% of a normal distribution with a mean and σ of the baseline mean SDF, Bonferroni corrected for multiple comparisons (where the number of independent observations was considered to be the duration of the epoch of interest divided by the width of the Gaussian kernel, 50ms). Similarly, a unit was considered to have significantly reduced firing within a given epoch if the mean ISI time series exceeded a threshold ISI value for at least 100ms. The threshold ISI value was defined as the upper 5% tail of a normal distribution with a mean and σ of the baseline mean ISI time series, Bonferroni corrected for multiple comparisons (where the number of independent observations was the mean number of ISIs within the epoch of interest).

### Speech onset- and cue- locking

For all units with significant changes in mean firing, we sought to determine whether the timing of these responses was more closely locked to the presentation of the cue or to the onset of the production, by examining the trial-to-trial relationship between RTs and neuronal response onsets. First, response onsets were estimated for individual trials. The trial-to-trial timing of an increase in firing was estimated by searching for bursts. The spiking pattern within each trial (after cue presentation) was examined to find a sequence of at least 3 spikes with the highest Poisson Surprise (PS) Burst index. For a given sequence of n spikes within time interval *T*, the PS Burst index was based on the probability of encountering n or more spikes within time interval *T*, given a Poisson-distributed spike train with a discharge rate *r*.

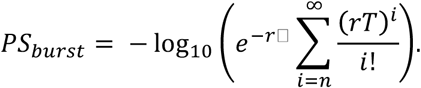

Similarly, the trial-to-trial timing of a decrease in firing was estimated by searching for pauses. The PS Pause index was based on the probability of encountering n or fewer spikes within time interval *T*, in a Poisson-distributed spike train:

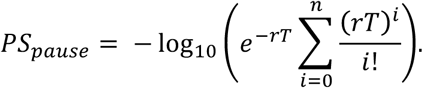

For both increase and decrease indices, *r* was estimated separately for each trial as the discharge rate across the entire trial, and T was the duration of the trial. Only trials with burst or pause sequences whose PS indices exceeded those found in that trial’s baseline epoch were considered for further analysis. For each trial, the onset time of the PS Burst (for units with significant excitatory responses) or PS Pause (for units with significant inhibitory responses) spike sequences was defined as the neuronal response increase or decrease onset, respectively.

Next, two intervals were correlated (Spearman rank correlation, MATLAB function corr) with the production latency across trials for each unit: 1. the interval between cue presentation and the neuronal response onset (neuronal response latency), and 2. the interval between the neuronal response onset and production onset (neuronal response to production interval). A unit’s response was considered to be temporally-locked to: 1. cue onset, if a significant change in activity during the cue epoch was observed, and the corresponding neuronal response to production interval was correlated (p<0.05) with production latency (Figure 6A-B), or 2. the onset of speech, if significant change in activity in the production epoch was observed, and the corresponding neuronal response latency was correlated (p<0.05) with production latency (Figure 6C-D). If both correlations were significant, then the unit’s response was considered to be both cue- and production -locked, i.e. its activity was temporally associated with both events.

**Figure 6.**
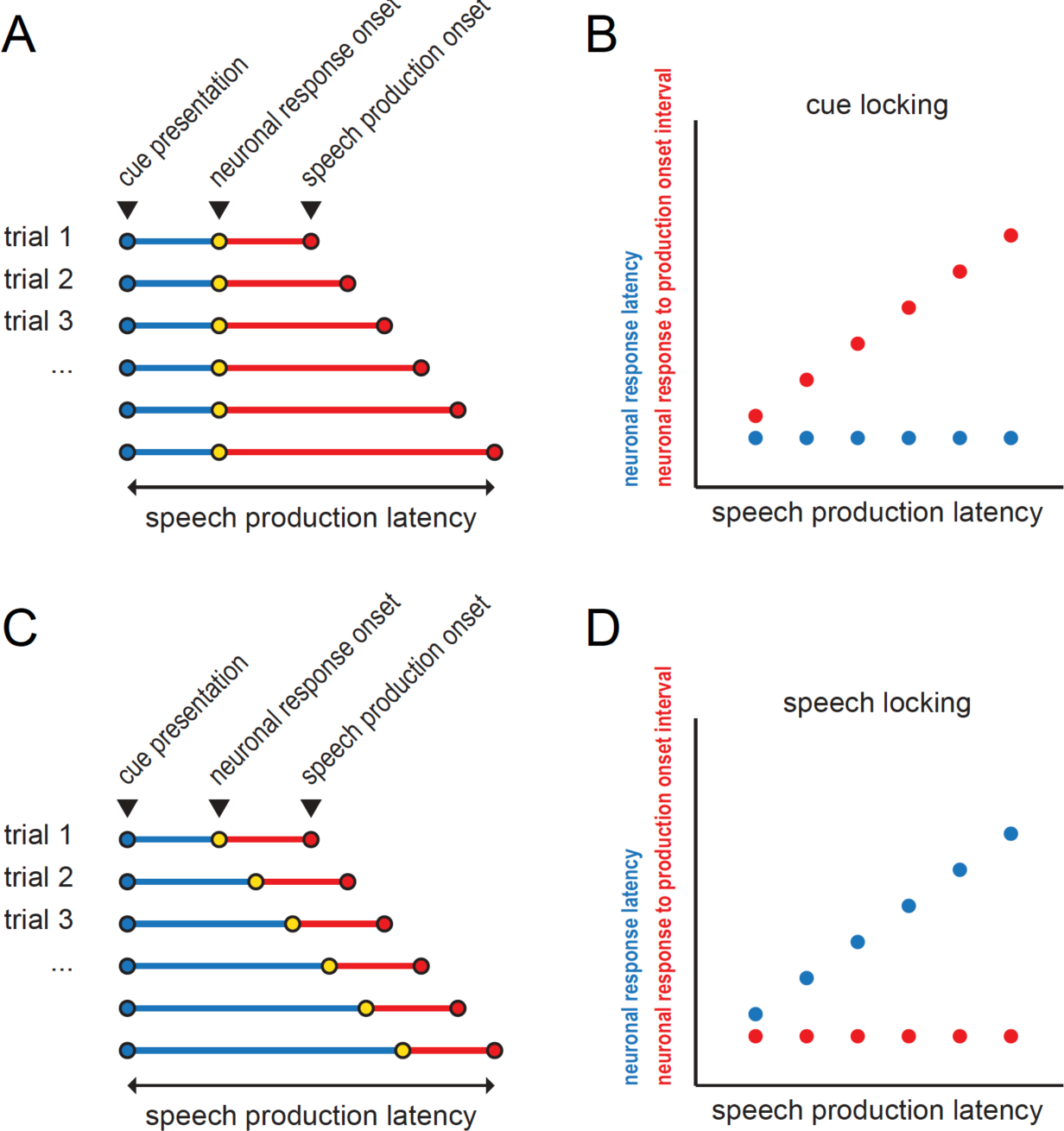
Schematic illustrating cue- and speech production-locking neuronal response types. (A) Hypothetical example of cue-aligned trials, illustrating a constant neuronal response latency with varying speech production latencies. (B) Corresponding correlation schematic showing that a significant correlation between neuronal response to production onset interval and speech production latency indicates cue-locking. (C) Hypothetical example of cue-aligned trials, illustrating a constant neuronal response to production onset interval with varying speech production latencies. (D) Corresponding correlation schematic showing that a significant correlation between the neuronal response latency and speech production latency indicates speech-locking.

### Anatomical localization of recording

Anatomical locations of microelectrode recordings were expressed in terms of the microelectrode recording-defined STN boundaries along each electrode trajectory. Thus, each microelectrode recording location was identified by its relative position within the Ben-Gun orientation (central, posterior or medial) and the percent depth through the STN within that trajectory (with 0% representing the ventral STN boundary and 100% representing the dorsal STN boundary).

---

**Supplementary Table 1.**
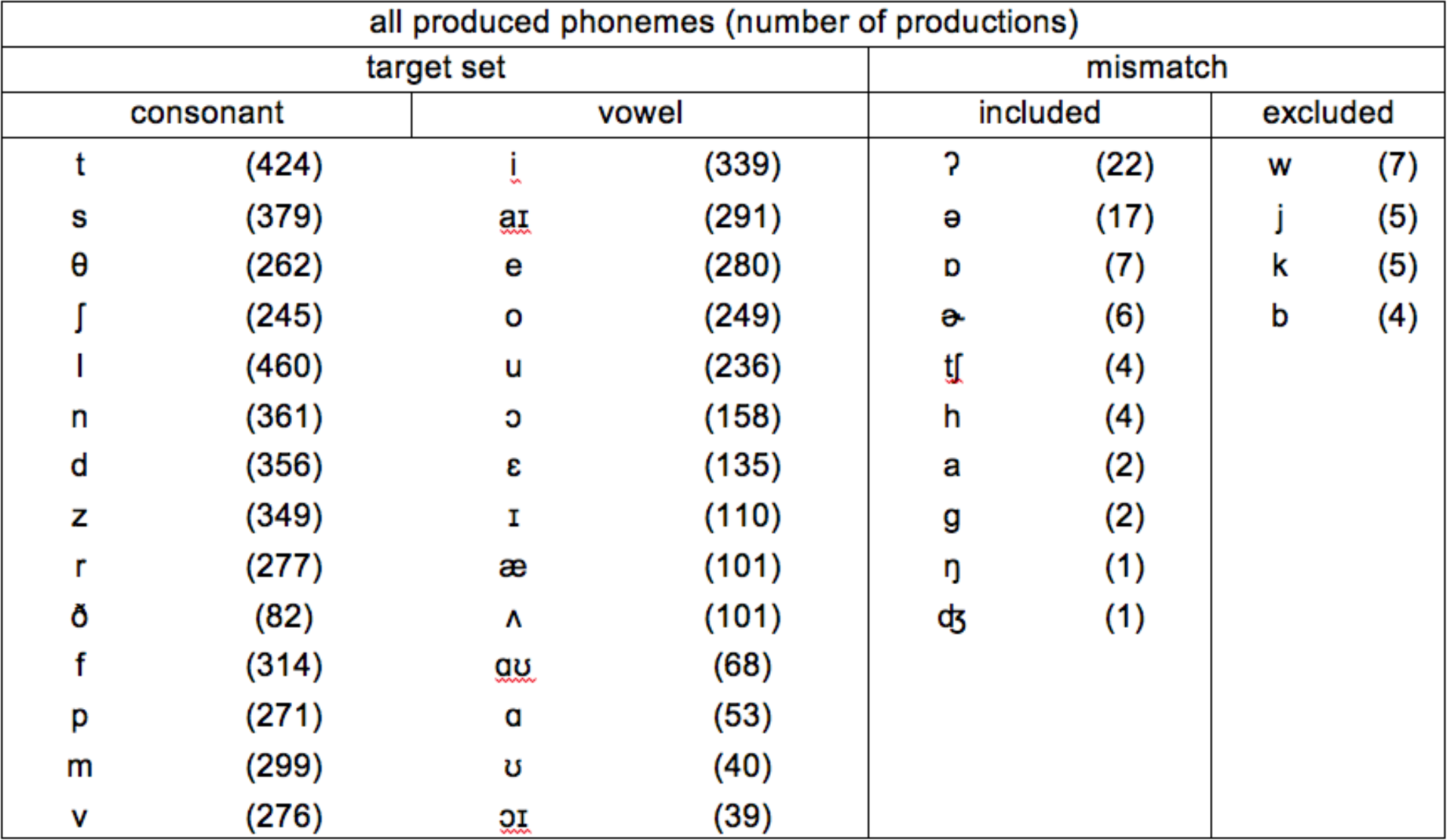
Phoneme frequency.

